# HCP5 prevents ubiquitination-mediated UTP3 degradation to inhibit apoptosis by activating c-Myc transcriptional activity

**DOI:** 10.1101/2022.06.13.495862

**Authors:** Yabing Nan, Qingyu Luo, Xiaowei Wu, Wan Chang, Pengfei Zhao, Shi Liu, Zhihua Liu

## Abstract

Inducing cancer cell apoptosis through cytotoxic reagents is the main therapeutic strategy for diverse cancer types. However, several antiapoptotic factors impede curative cancer therapy by driving cancer cells to resist cytotoxic agent-induced apoptosis, thus leading to refractoriness and relapse. To define critical antiapoptotic factors that contribute to chemoresistance in esophageal squamous cell carcinoma (ESCC), we generated two pairs of parental and apoptosis-resistant cell models through cisplatin (DDP) induction and then performed whole-transcriptome sequencing. We identified the long noncoding RNA (lncRNA) histocompatibility leukocyte antigen complex P5 (HCP5) as the chief culprit for chemoresistance. Mechanistically, HCP5 interacts with UTP3 small subunit processome component (UTP3) and prevents UTP3 degradation from E3 ligase tripartite motif containing 29 (TRIM29)-mediated ubiquitination. UTP3 then recruits c-Myc to activate vesicle-associated membrane protein 3 (VAMP3) expression. Activated VAMP3 suppresses caspase-dependent apoptosis and eventually leads to chemoresistance. Accordingly, the expression level of the HCP5/UTP3/c-Myc/VAMP3 axis in chemoresistant patients is significantly higher than that in chemosensitive patients. Thus, our study demonstrated that the HCP5/UTP3/c-Myc/VAMP3 axis plays an important role in the inhibition of cancer cell apoptosis and that HCP5 can be a promising chemosensitive target for cancer treatment.

## Introduction

Apoptosis is the most frequent type of regulated cell death and is involved in a variety of biological processes, while resistance to apoptosis is a hallmark of cancer that contributes to treatment failure (1–3). Several targeted inhibitors of apoptosis molecules, such as B-cell lymphoma 2 inhibitors (4) and myeloid cell leukemia 1 (MCL1) inhibitors (5), have shown promising effects in clinical applications. However, these inhibitors are applicable only to specific cancer subtypes and produce tolerance during use, resulting in ineffective treatment of cancer patients (6). Therefore, identifying new proapoptotic and antiapoptotic molecules and revealing their functional mechanisms are important for the establishment of new anticancer strategies.

Esophageal cancer is a deadly malignancy that ranks sixth in the cause of overall cancer mortality (7). Esophageal squamous cell carcinoma (ESCC), the main subtype of esophageal cancer, accounts for approximately 90% of all cases (8). Multidisciplinary treatment has improved the overall survival of patients with esophageal cancer compared to surgery alone. Neoadjuvant chemoradiotherapy is the standard of care for the treatment of patients with locally advanced ESCC (9). Cisplatin (DDP)- and 5-fluorouracil-based chemotherapy regimens have been widely adopted as neoadjuvant, radical, and palliative therapies for esophageal cancer over several decades (10,11). However, patients with esophageal cancer usually have a poor prognosis owing to therapeutic resistance. Since the dysregulation of the cell apoptosis pathway is critical for resistance to chemotherapy (12), it is essential to elucidate the detailed mechanisms by which cancer cells develop apoptosis resistance induced by cytotoxic chemotherapy.

Long noncoding RNAs (lncRNAs) participate in transcriptional and posttranscriptional gene regulation. At the transcriptional level, a model has been established in which lncRNAs bridge DNA and proteins by enhancing or blocking the recruitments of transcription regulators (13). Additional mechanisms by which lncRNAs regulate gene transcription include acting as RNA sponges or competitive endogenous RNAs (ceRNAs) through crosstalk and competition (14,15). LncRNAs can also affect the protein stability of transcriptional regulators via the ubiquitin–proteasome system (UPS) (16,17). Histocompatibility leukocyte antigen complex P5 (HCP5) is a lncRNA initially identified in 1993 (18). Recently, the expression of HCP5 has been reported to be upregulated in multiple cancers and is associated with malignant phenotypes such as tumor growth, metastasis, chemoresistance and cancer stemness (19–21). However, the underlying mechanism by which HCP5 mediates cancer cell apoptosis and gene transcriptional regulation remains elusive.

In this study, by performing whole-transcriptome sequencing of two pairs of DDP-resistant and parental cell models (22), we identified HCP5 as an antiapoptotic lncRNA, which is the most significantly upregulated lncRNA in DDP-resistant cells compared with parental cells. We found that HCP5 interacts with and stabilizes UTP3 small subunit processome component (UTP3) by blocking E3 ligase tripartite motif containing 29 (TRIM29)-mediated degradation. Elevated UTP3 then activates VAMP3 transcription by recruiting c-Myc to the promoter region of VAMP3. Thus, our study revealed a previously unknown mechanism by which the lncRNA HCP5 affects UTP3 protein stability and subsequent c-Myc transcriptional activities during chemoresistance.

## Results

### HCP5 augments chemoresistance and correlates with poor prognosis in ESCC

To define critical ncRNA that contributes to chemoresistance in ESCC, we established two DDP-resistant cell models, KYSE450/DDP and YES2/DDP, and then performed whole-transcriptome sequencing of resistant and parental cells (22). We identified 311 and 352 significantly upregulated ncRNAs in the two DDP-resistant cell models, respectively, and HCP5 was the most significantly upregulated ncRNA (Fig. 1A). The striking elevation of HCP5 in chemoresistant cells was confirmed by an RT–qPCR assay in an independent culture of DDP-resistant and parental cells (Fig. 1B). Interestingly, HCP5 expression was also significantly elevated after short-term induction with DDP in a time-dependent manner (6, 12, or 24 h) (Fig. 1C). These results suggested that elevated HCP5 expression could serve as both a fast response by cancer cells to survive from cytotoxic pressure and a long-term mechanism contributing to acquired chemoresistance. To assess the potential roles of HCP5 in clinical outcomes, we first examined HCP5 expression in 215 ESCC patient samples by performing in situ hybridization (ISH). The results revealed that high HCP5 expression in ESCC patients significantly correlated with poor prognosis (Fig. 1D). In addition, we analyzed the HCP5 expression levels in 168 pairs of ESCC and normal esophageal epithelial tissues and found that HCP5 expression was increased in ESCC tissues compared with normal tissues (Fig. 1E). Consistent with our ESCC cohort results, an analysis of 53 pairs of ESCC and normal tissues in the Gene Expression Omnibus (GEO) database showed that HCP5 exhibited higher expression in the cancer tissues than in the normal counterparts (Fig. 1F). No significant correlation was found between the HCP5 expression level and tumor depth, lymph node metastasis, distant metastasis or tumor grade (Supplementary Fig. S1A).

**Fig. 1.**
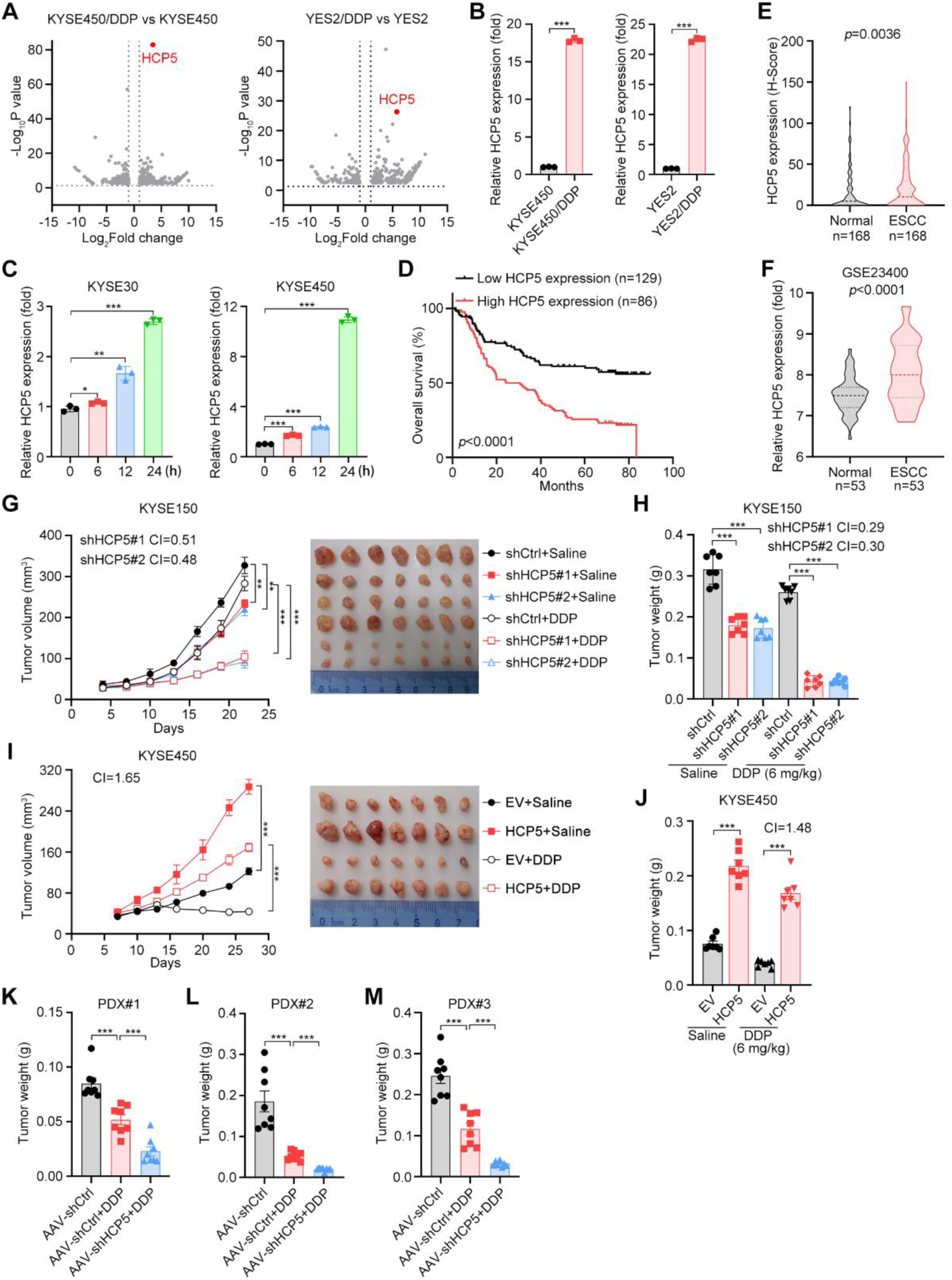
HCP5 augments chemoresistance and correlates with poor prognosis in ESCC. **A** Volcano plots showing upregulated and downregulated ncRNAs in KYSE450/DDP and YES2/DDP cells compared to their parental cells. **B** RT–qPCR detection of HCP5 expression in the parental and DDP-resistant cell models. The data are presented as the means ± s.d.; two-tailed *t* test, \*\*\**P* < 0.001; n = 3. **C** RT–qPCR detection of HCP5 expression in KYSE30 and KYSE450 cells treated with DDP for 0, 6, 12, and 24 h. The data are presented as the means ± s.d.; two-tailed *t* test, \*\*\**P* < 0.001, \*\**P* < 0.01, \**P* < 0.05; n = 3. **D** Overall survival of ESCC patients with low and high HCP5 expression levels (divided by the mean expression value). Kaplan–Meier survival plots are shown. **E** Statistical analysis of HCP5 expression in 168 paired ESCC tissues and adjacent normal tissues. Box plot representation: from top to bottom—maximum, 75th percentile; median, 25th percentile; and minimum values; two-tailed *t* test. **F** Statistical analysis of HCP5 expression in 53 paired ESCC tissues and adjacent normal tissues in the GEO database. Box plot representation: from top to bottom—maximum, 75th percentile; median, 25th percentile; and minimum values; two-tailed *t* test. **G-J** Tumor volumes and representative images (G and I) and tumor weights (H and J) of the xenografts derived from the indicated cells. The data are presented as the means ± s.e.m.; two-tailed *t* test, \*\*\**P* < 0.001, \*\**P* < 0.01; n = 7. **K-M** Tumor weights in the three PDX models from ESCC patients treated with AAV-shCtrl or AAV-shHCP5. The data are presented as the means ± s.e.m.; two-tailed *t* test, \*\*\**P* < 0.001; n = 8.

To investigate the functions of HCP5 in the progression of ESCC, we used lentiviral short hairpin RNA (shRNA) to knockdown HCP5 expression in the KYSE150 cell line, which showed relatively high HCP5 expression (Supplementary Fig. S1B). A stable HCP5 overexpression model was also established with KYSE450 cells that have low endogenous HCP5 expression levels (Supplementary Fig. S1B). In vitro proliferation experiments indicated that knocking down HCP5 expression markedly decreased the growth rates of ESCC cells, and overexpression of HCP5 promoted cell proliferation (Supplementary Fig. S1C). Cell viability assays showed that HCP5 depletion dramatically increased cell sensitivity to DDP, and exogenous expression of HCP5 promoted chemoresistance (Supplementary Fig. S1D). We next studied the effects of HCP5 on ESCC progression in vivo. Xenograft assays indicated that HCP5 silencing sensitized KYSE150 cells to DDP, with a combination index (CI) from 0.29 to 0.51 for tumor volume and weight (Fig. 1G, H). On the other hand, HCP5 overexpression in KYSE450 cells promoted their resistance to DDP treatment, with CIs of 1.65 and 1.48 for tumor volume and weight, respectively (Fig. 1I, J). To further define the potential of targeting HCP5 to increase chemosensitivity, we established three patient-derived xenograft (PDX) models of ESCC and then inhibited HCP5 by adeno-associated virus (AAV)-mediated administration of shRNAs. In all three PDX models, in vivo administration of AAV-shHCP5 markedly sensitized ESCC samples to DDP treatment (Fig. 1K-M and Supplementary Fig. S1E-G).

### HCP5 impedes caspase-dependent apoptotic signaling

To dissect the molecular mechanism of how HCP5 contributes to chemoresistance, we performed RNA sequencing (RNA-seq) with the HCP5-silenced KYSE150 and KYSE510 cell lines. Compared to control cells, HCP5-silenced cells showed significant enrichment of the apoptosis pathway from the gene set enrichment analysis (GSEA) (Fig. 2A). Consistently, caspase 3/7 activity assays showed that the silencing of HCP5 dramatically increased the activation of apoptotic signaling cascades upon DDP treatment (Fig. 2B, C). On the other hand, overexpression of HCP5 significantly suppressed caspase activation under cytotoxic exposure (Fig. 2D, E). The effect of HCP5 on the caspase-dependent apoptosis pathway was further confirmed by the detection of apoptotic markers (cleaved caspase-3, cleaved caspase-7 and cleaved PARP) by western blotting (Fig. 2F-I).

**Fig. 2.**
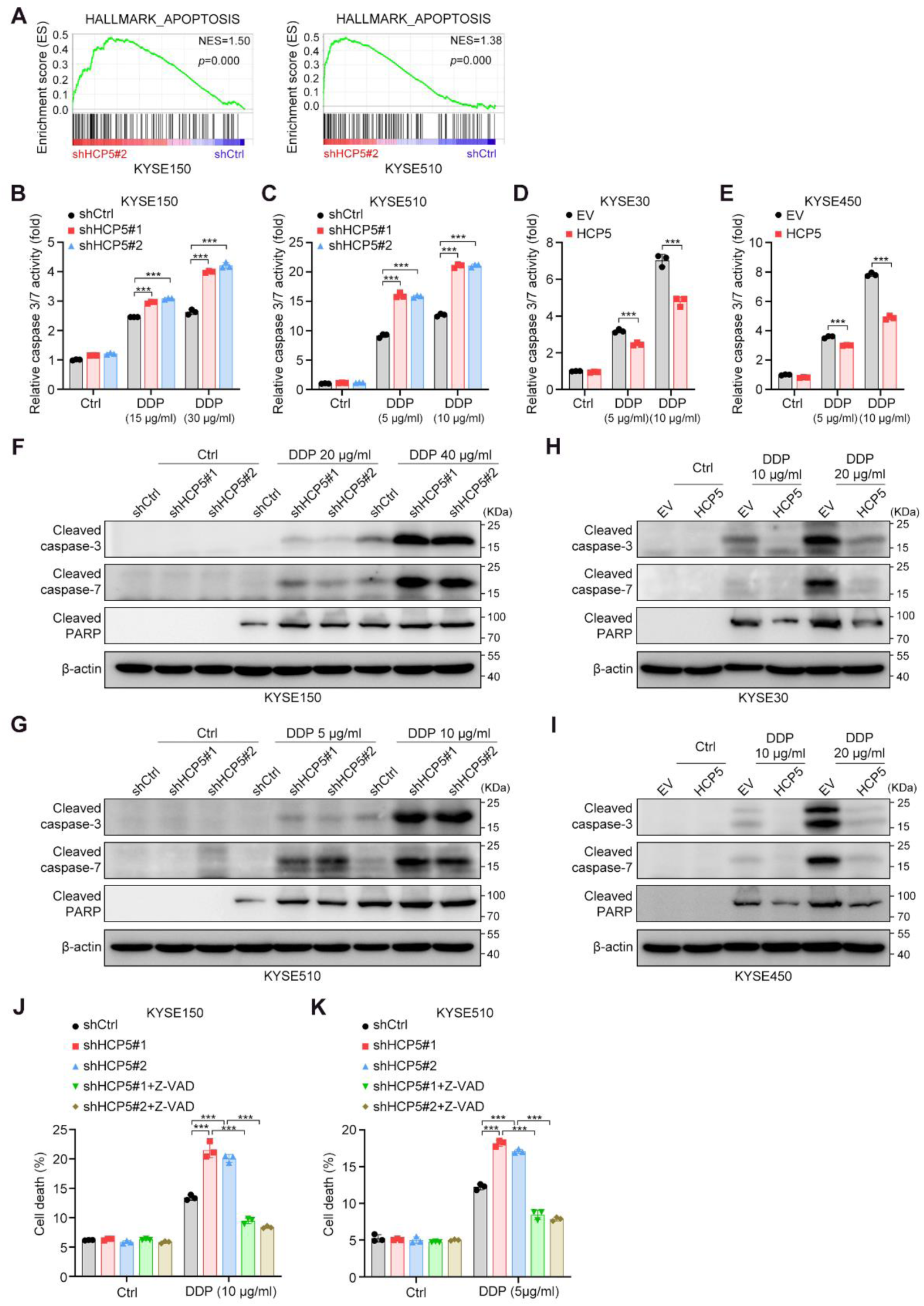
HCP5 impedes caspase-dependent apoptotic signaling. **A** Pathways enriched by GSEA based on RNA-seq data obtained with HCP5-silenced and control KYSE150 and KYSE510 cells. **B-E** Detection of caspase 3/7 activity in the indicated cells. The data are presented as the means ± s.d.; two-tailed *t* test, \*\*\**P* < 0.001; n = 3. **F, G** Western blot analyses of the expression of cleaved caspase-3, cleaved caspase-7 and cleaved PARP in control and HCP5-silenced KYSE150 (F) or KYSE510 (G) cells. Cells were exposed to the indicated drug concentrations for 24 h before detection. **H, I** Western blot analyses of the expression of cleaved caspase-3, cleaved caspase-7 and cleaved PARP in KYSE30 (H) or KYSE450 (I) cells expressing EV or HCP5. Cells were exposed to the indicated drug concentrations for 24 h before detection. **J, K** Cell death fractions of the control and HCP5-silenced KYSE150 (J) and KYSE510 (K) cells treated with DDP with or without Z-VAD treatment (50 μM). The data are presented as the means ± s.d.; two-tailed *t* test, \*\*\**P* < 0.001; n = 3.

Since apoptosis can also be activated through caspase-independent mechanisms, we treated HCP5-silenced KYSE150 and KYSE510 cells with the pancaspase inhibitor Z-VAD-FMK and then performed annexin V-FITC/propidium iodide (PI) double staining assays. The results indicated that silencing HCP5 enhanced the apoptosis rate of the KYSE150 and KYSE510 cells and promoted their sensitivity to DDP treatment, while treatment with Z-VAD-FMK completely abrogated the proapoptotic function of HCP5 silencing with or without DDP treatment (Fig. 2J, K and Supplementary Fig. S2A, B). In conclusion, these results indicated that HCP5 impedes caspase-dependent apoptotic signaling.

### HCP5 directly interacts with and stabilizes UTP3 by preventing its ubiquitination

To further explore the mechanism underlying HCP5 impeding apoptosis and subsequently contributing to chemoresistance, we first assessed the subcellular localization of HCP5 in ESCC cells. An RNA fluorescence ISH (FISH) assay showed that HCP5 was distributed in both the nucleus and cytoplasm (Fig. 3A), which was confirmed by nuclear and cytoplasmic RNA fractionation analysis using RT–qPCR (Supplementary Fig. S3A). These results suggest that in addition to previously reported functioning as a miRNA sponge (23–25), HCP5 may also participate in transcriptional regulation by interacting with transcriptional regulators in the nucleus. To validate this hypothesis, we determined HCP5-interacting proteins by performing RNA pull-down assays followed by mass spectrometry (MS). In total, 77 and 113 HCP5-specific interacting proteins were identified in KYSE150 and KYSE510 cells, respectively, and 8 of these proteins were consistently identified in both cell lines (Fig. 3B). Among the 8 common proteins, we primarily focused on UTP3 due to its potential in gene transcriptional regulation. UTP3 was identified to derepress silenced mating-type genes in yeast (26), while its detailed functioning mechanisms, especially in the human context, remain unclear.

**Fig. 3.**
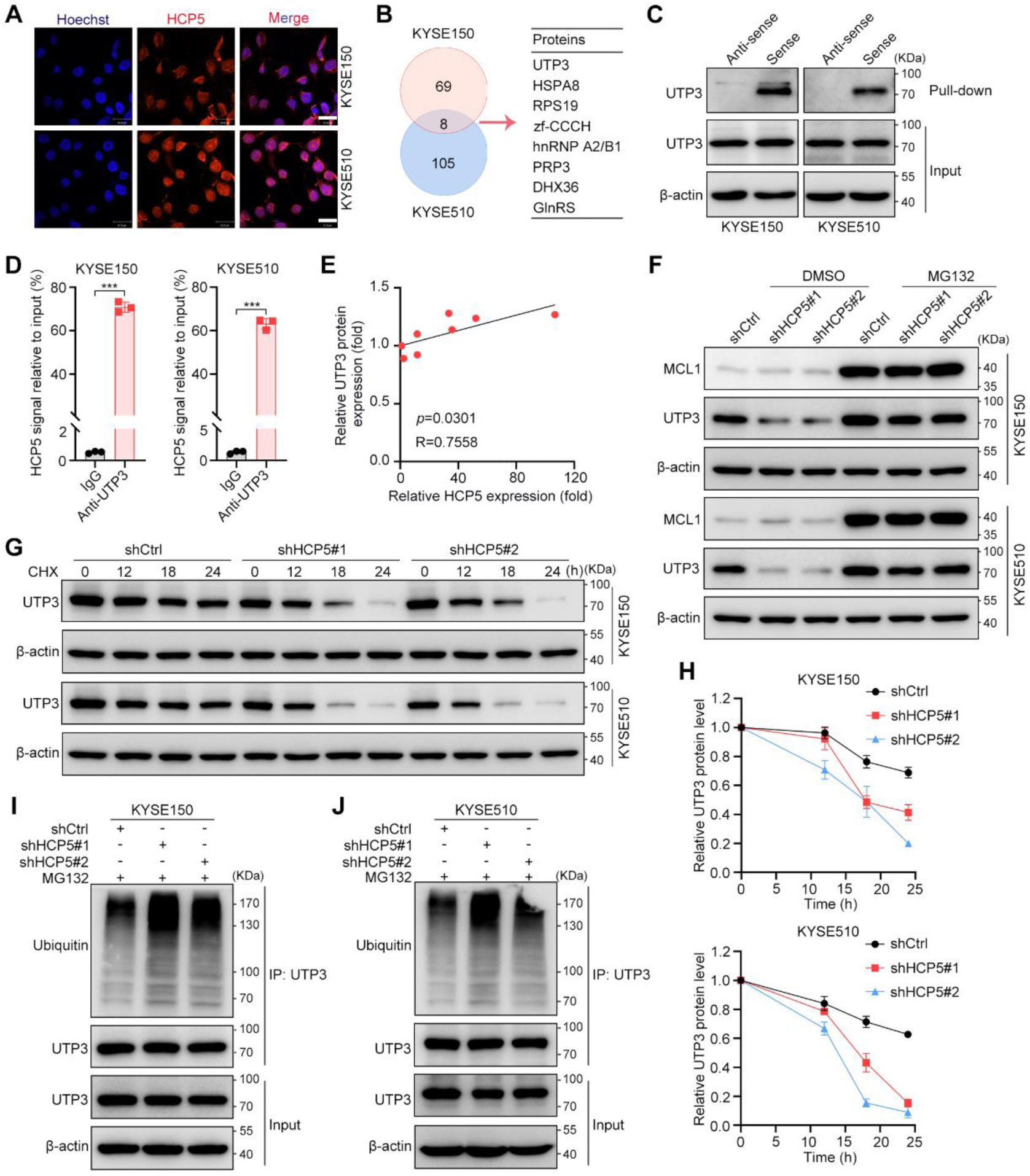
HCP5 interacts with and stabilizes UTP3 by preventing its ubiquitination. **A** FISH detection of HCP5 localization in KYSE150 and KYSE510 cells. Nuclei were stained with Hoechst 33342. Scale bar, 30 μm. **B** List of HCP5-interacting proteins identified using MS with KYSE150 and KYSE510 cells. **C** Western blot analyses of UTP3 in precipitates from RNA pull-down assays performed using antisense or sense sequences of HCP5. **D** RT–qPCR measurement of the HCP5 level in precipitates from RIP assays performed using IgG or anti-UTP3 antibodies. The data are presented as the means ± s.d.; two-tailed *t* test, \*\*\**P* < 0.001; n = 3. **E** Correlation analysis of HCP5 and UTP3 protein expression in ESCC cell lines. Spearman correlation coefficients are shown. **F** Western blot analyses of UTP3 and MCL1 expression in control and HCP5-silenced KYSE150 and KYSE510 cells treated with DMSO or MG132 for 8 h. **G** Western blot analyses of UTP3 expression in control and HCP5-silenced KYSE150 and KYSE510 cells subjected to a CHX pulse-chase assay. **H** Quantitative analyses of UTP3 expression in control and HCP5-silenced KYSE150 and KYSE510 cells subjected to the CHX pulse-chase assay. Quantification of UTP3 expression relative to *β*-actin expression is shown. The data are reported as the means ± s.e.m.; n = 3. **I, J** Western blot detection to assess UTP3 ubiquitination in control and HCP5-silenced KYSE150 (I) and KYSE510 cells (J).

To verify the specific interaction between HCP5 and UTP3, we performed independent RNA pull-down assays and then detected UTP3 by western blotting. The results indicated that the HCP5 sense strand, but not the antisense strand, interacted with UTP3 (Fig. 3C). The interaction between HCP5 and UTP3 was also confirmed by RNA immunoprecipitation (RIP) assays (Fig. 3D). Next, we determined whether HCP5 affects UTP3 expression levels. Interestingly, although the mRNA levels of UPT3 remained unchanged upon either HCP5 knockdown or overexpression (Supplementary Fig. S3B, C), the UTP3 protein levels were markedly affected by HCP5 (Supplementary Fig. S3D). We then examined correlations between HCP5 and UTP3 mRNA and protein levels by RT–qPCR and western blotting assays, respectively. As expected, HCP5 expression was positively correlated with UTP3 protein expression in ESCC cell lines (Fig. 3E and Supplementary Fig. S3E). On the other hand, there was no correlation between HCP5 and UTP3 mRNA expressions (Supplementary Fig. S3E, F).

As a crucial posttranslational modification process, ubiquitination plays a critical role in regulating the stability and function of proteins (27,28); thus, we hypothesized that HCP5 stabilized UTP3 by affecting UPS-mediated protein degradation. To validate our hypothesis, we treated control and HCP5-silenced KYSE150 and KYSE510 cells with the proteasome inhibitor MG132 and found that HCP5 silencing-mediated protein degradation of UTP3 was reestablished by MG132 but not DMSO (Fig. 3F). In addition, MG132 increased UTP3 protein levels and abolished the effect of HCP5 overexpression on UTP3 protein expression (Supplementary Fig. S3G). MCL1 was included as a positive control based on a previous report (29) (Fig. 3F and Supplementary Fig. S3G). Cycloheximide (CHX) chase assays showed that silencing HCP5 expression resulted in a shortened half-life of the endogenous UTP3 protein (Fig. 3G, H), whereas overexpression of HCP5 led to a prolonged half-life of the UTP3 protein (Supplementary Fig. S3H, I). Finally, we performed immunoprecipitation (IP) assays with control and HCP5-silenced KYSE150 and KYSE510 cells using an anti-UTP3 antibody and detected ubiquitinated protein levels. The results indicated that HCP5 silencing significantly increased the levels of ubiquitinated UTP3 (Fig. 3I, J). Taken together, these results indicate that HCP5 stabilizes UTP3 protein levels by inhibiting its degradation by the UPS.

### HCP5 stabilizes UTP3 by blocking the interaction between UTP3 and TRIM29

To ask how HCP5 prevents ubiquitination of UTP3, we hypothesized that HCP5 could prevent the interaction between E3 ligases and UTP3. Thus, we performed IP assays followed by MS to identify the specific E3 ligase that mediates UTP3 protein degradation. A total of 6 E3 ligases were identified to interact with UTP3 in both KYSE150 and KYSE510 cell lines (Fig. 4A). We next designed shRNA sequences to knockdown these 6 E3 ligases separately in KYSE150 and KYSE510 cells and then detected the UTP3 expression levels. The results indicated that silencing of none of these six E3 ligases had an obvious effect on UTP3 mRNA levels (Supplementary Fig. S4A, B). On the other hand, the UTP3 protein level was remarkably upregulated in both cell lines after TRIM29 knockdown but not in the context of silencing the other five E3 ligases (Fig. 4B and Supplementary Fig. S4C). Moreover, the UTP3 protein level, but not its mRNA level, was gradually reduced after exogenous expression of an increasing amount of TRIM29 (Fig. 4C and Supplementary Fig. S4D).

**Fig. 4.**
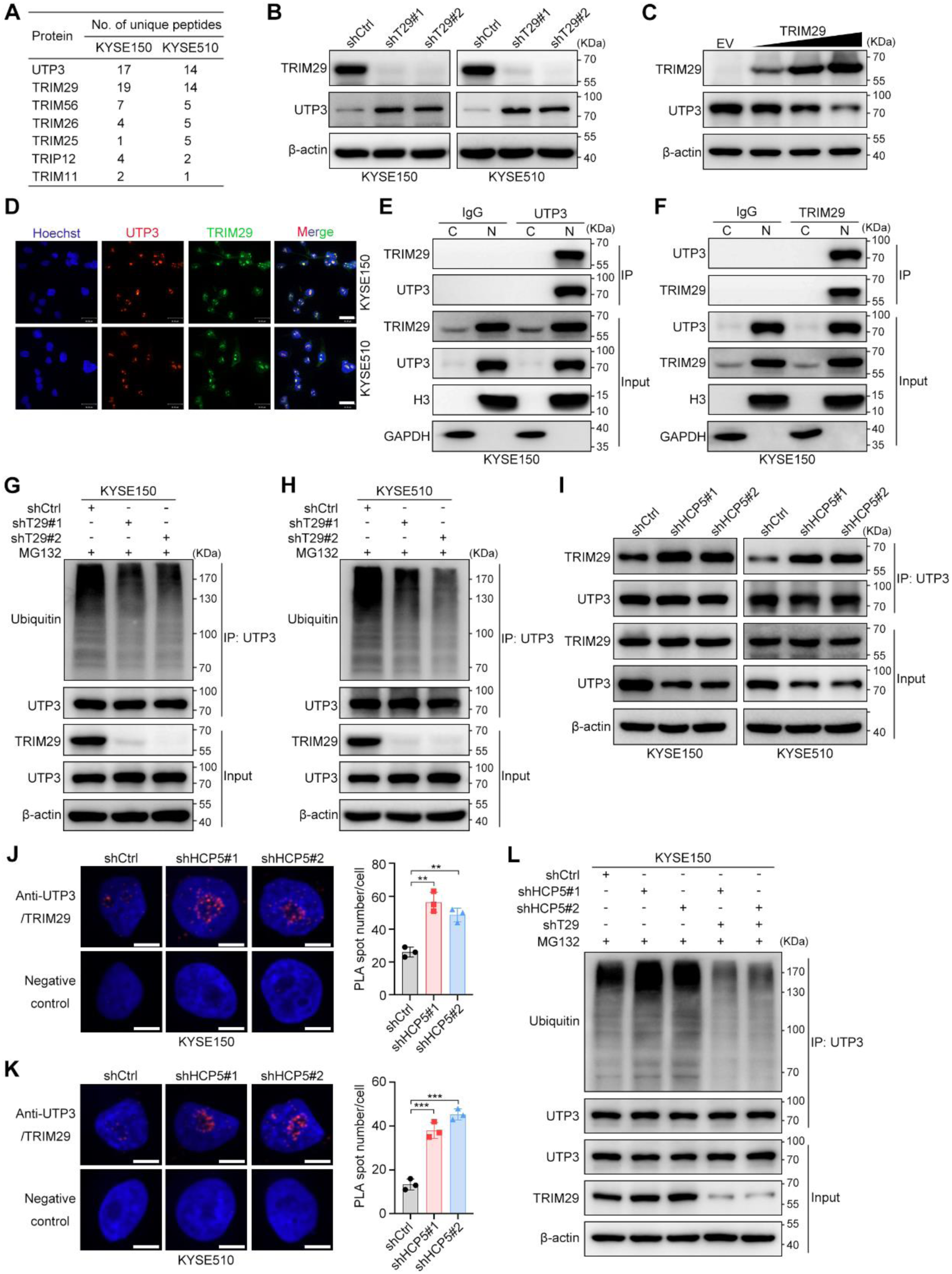
HCP5 stabilizes UTP3 by blocking the interaction between UTP3 and TRIM29. **A** List of UTP3-interacting E3 ligases identified using MS in KYSE150 and KYSE510 cells. **B** Western blot analyses of TRIM29 and UTP3 expression in control and TRIM29-silenced KYSE150 and KYSE510 cells. **C** Western blot analyses of TRIM29 and UTP3 expression in 293T cells transfected with increasing amounts of TRIM29 (0.5 μg, 1.0 μg and 2.0 μg). **D** Confocal microscopy assessment of UTP3 and TRIM29 colocalization in the indicated cells. Nuclei were stained with Hoechst 33342. Scale bars, 30 µm. **E, F** Western blot analyses of TRIM29 and UTP3 distribution after co-IP assays with the indicated cells (C, cytoplasm; N, nucleus). **G, H** Western blot analyses to assess UTP3 ubiquitination in control and TRIM29-silenced KYSE150 (G) and KYSE510 cells (H). **I** Western blot analyses of TRIM29 and UTP3 in co-IP assays performed with control and HCP5-silenced KYSE150 and KYSE510 cells. **J, K** Confocal microscopy assessment of PLA spots (red) in the indicated cells. Each spot indicates a single interaction between UTP3 and TRIM29 proteins. Nuclei were stained with DAPI (blue). Scale bars, 10 µm. The data are presented as the means ± s.d.; two-tailed *t* test, \*\*\**P* < 0.001, \*\**P* < 0.01; n = 3. **L** Western blot analyses to assess UTP3 ubiquitination in the indicated cells.

TRIM29 is a new member of the TRIM family (30). TRIM29-mediated ubiquitination often leads to protein degradation and has been reported to exert important functions in several human cancer types (31–33). However, how interactions with ncRNAs affect TRIM29-mediated target protein degradation remains unknown. Therefore, we next investigated the effect of TRIM29 on HCP5-mediated UTP3 protein stabilization. Double-labeling immunofluorescence showed that UTP3/TRIM29 exhibited extensive colocalization in the nucleus (Fig. 4D). The specific interaction between TRIM29 and UTP3 in the nucleus was further validated by co-IP assays performed with nuclear and cytoplasmic protein lysates separately (Fig. 4E, F). Finally, ubiquitination assays showed that TRIM29 knockdown markedly decreased the levels of ubiquitinated UTP3 in ESCC cells (Fig. 4G, H), confirming that TRIM29 is a bona fide E3 ligase of UTP3.

We next determined whether HCP5 stabilized UTP3 by affecting TRIM29-mediated ubiquitination. Neither HCP5 knockdown nor overexpression affected TRIM29 protein expression (Supplementary Fig. S4E, F). Moreover, no direct interaction between HCP5 and TRIM29 was found in the RNA pull-down assays (Supplementary Fig. S4G). Interestingly, the silencing of HCP5 in KYSE150 and KYSE510 cells promoted the interaction between TRIM29 and UTP3, as indicated by co-IP assays (Fig. 4I) and proximity ligation (PLA) assays (Fig. 4J, K). These results indicate that HCP5 may stabilize UTP3 by interfering with the interaction between TRIM29 and UTP3. Supporting this hypothesis, ubiquitination assays showed that the silencing of TRIM29 completely abrogated the increased ubiquitination of UTP3 upon HCP5 depletion (Fig. 4L).

### UTP3 mediates the antiapoptotic and chemoresistant roles of HCP5

We next asked whether UTP3 mediates the antiapoptotic and chemoresistant phenotypes triggered by HCP5. Cell viability assays indicated that the overexpression of UTP3 restored cell viability that had been diminished by HCP5 silencing (Supplementary Fig. S5A). On the other hand, knocking down UTP3 expression abolished HCP5 overexpression-mediated increases in cell viability (Supplementary Fig. S5B). The critical role of UTP3 in HCP5-mediated chemoresistance was also confirmed by in vivo xenograft assays (Fig. 5A-D). To further study whether UTP3 mediates the function of HCP5 in inhibiting apoptotic signaling, rescue assays were performed, and caspase activation was detected. As expected, Caspase 3/7 activity assays demonstrated that UTP3 mediated the antiapoptotic function of HCP5 upon DDP treatment (Fig. 5E, F). The same conclusion was obtained by western blotting for apoptotic markers (Fig. 5G, H). Accordingly, the proapoptotic functions of UTP3 silencing were completely abrogated after treatment with the pancaspase inhibitor Z-VAD-FMK (Fig. 5I and Supplementary Fig. S5C).

**Fig. 5.**
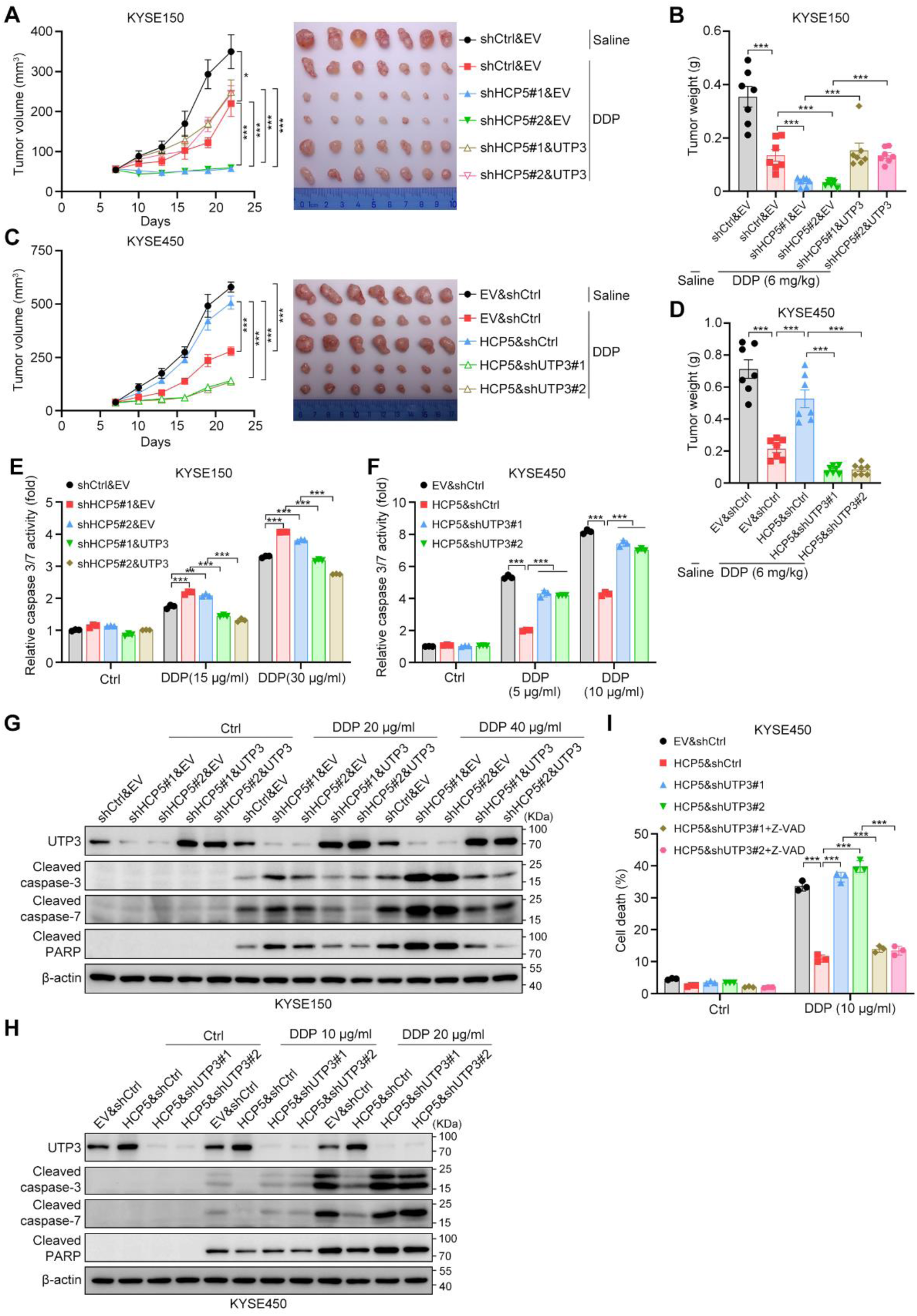
HCP5/UTP3 inhibits caspase-dependent cell apoptosis. **A-D** Tumor volumes and representative images (A and C) and weights (B and D) of xenograft tumors derived from the indicated cells. The data are presented as the mean ± s.e.m.; two-tailed t test, \*\*\**P* < 0.001, \**P* < 0.05; n = 7. **E, F** Detection of caspase 3/7 activity in the indicated cells. The data are presented as the means ± s.d.; two-tailed *t* test, \*\*\**P* < 0.001, \*\**P* < 0.01; n = 3. **G, H** Western blot analyses of the expression of UTP3, cleaved caspase-3, cleaved caspase-7 and cleaved PARP in the indicated cells. The cells were exposed to the indicated drug concentrations for 24 h before detection. **I** Cell death fractions of the indicated cells treated with DDP with or without Z-VAD (50 μM). The data are presented as the means ± s.d.; two-tailed *t* test, \*\*\**P* < 0.001; n = 3.

### HCP5 activates VAMP3 at transcriptional level

To investigate the potential mechanism by which UTP3 mediates HCP5 inhibition of cell apoptosis, we performed assay for transposase-accessible chromatin sequencing (ATAC-seq) and RNA-seq with control and HCP5-silenced KYSE150 and KYSE510 cells. A total of 8 genes (including HCP5) were consistently downregulated in HCP5-silenced KYSE150 and KYSE510 cells (Supplementary Fig. S6A). Among those 7 downregulated genes except for HCP5, NUS1P1 and BCLAF1P1 are pseudogenes, so we selected the other five genes (VAMP3, NEK7, NUS1, TMED10 and SLC16A14) for further validation. The ATAC-seq results indicated that only VAMP3 showed an obvious loss of peak in HCP5-depleted cells (Fig. 6A-C), indicating that VAMP3 may be a direct target gene of HCP5.

**Fig. 6.**
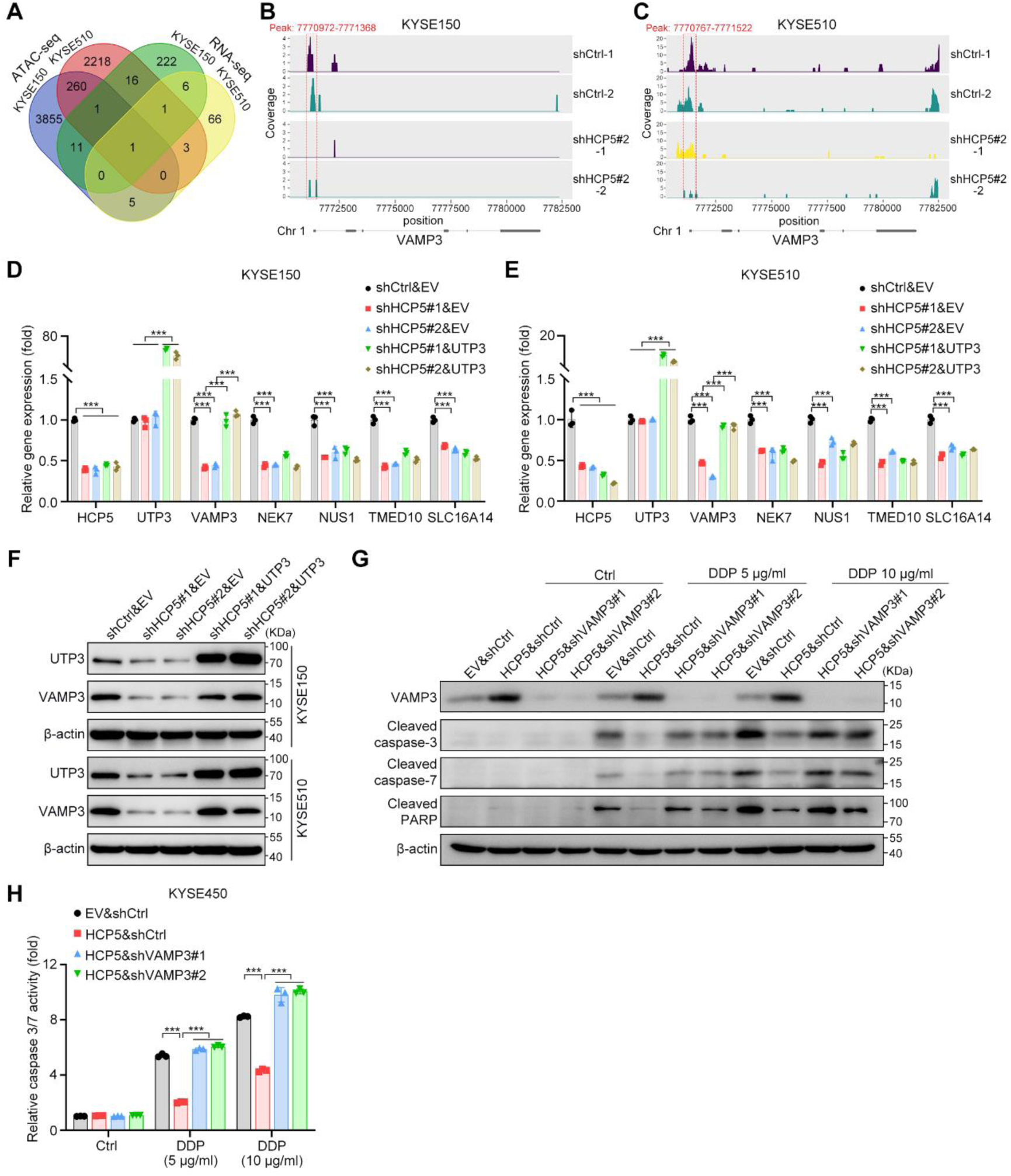
HCP5 activates VAMP3 expression at transcriptional level. **A** Number of differently expressed genes in HCP5-silenced KYSE150 and KYSE510 cells compared to control cells, as determined with ATAC-seq and RNA-seq. **B, C** Unique peaks identified at VAMP3 promotor region in the indicated cells as determined with ATAC-seq. **D, E** RT–qPCR measurement of gene expressions in the indicated cells. The data are presented as the means ± s.d.; two-tailed t test, \*\*\**P* < 0.001; n = 3. **F** Western blot analyses of UTP3 and VAMP3 expressions in the indicated cells. **G** Western blot analyses of the expression of VAMP3, cleaved caspase-3, cleaved caspase-7 and cleaved PARP in the indicated cells. The cells were exposed to the indicated drug concentrations for 24 h before detection. **H** Detection of caspase 3/7 activity in the indicated cells. The data are presented as the means ± s.d.; two-tailed *t* test, \*\*\**P* < 0.001; n = 3.

We next asked whether UTP3 transcriptionally regulates VAMP3 expression. RT–qPCR confirmed that overexpression of UTP3 completely rescued the downregulation of VAMP3 mRNA expression levels that had been mediated by HCP5 knockdown in KYSE150 and KYSE510 cells; however, the expression levels of the other four genes were not rescued (Fig. 6D, E). Accordingly, western blotting showed that UTP3 overexpression rescued the reduced VAMP3 protein expression in HCP5-depleted cells (Fig. 6F). Finally, we validated the functional relationships among HCP5, UTP3 and VAMP3 regarding apoptosis and chemoresistance. The results showed that knocking down VAMP3 expression abrogated the inhibitory effect of HCP5 on the apoptosis of cells treated with DDP, as indicated by both western blotting (Fig. 6G) and caspase 3/7 activity assays (Fig. 6H). Taken together, these results indicate that HCP5/UTP3 transcriptionally activate VAMP3 to inhibit cell apoptosis.

### UTP3 recruits c-Myc to transcriptionally activate VAMP3

The GSEA of RNA-seq data showed significant enrichment of MYC targets with HCP5 (Fig. 7A, B), indicating that c-Myc-mediated transcriptional activity may contribute to the phenotypes underlying the HCP5/UTP3/VAMP3 axis. To validate this hypothesis, we first generated a sequence logo of MYC by the JASPAR database (34) (Fig. 7C), and further analysis revealed that MYC could bind several sequences in the VAMP3 promoter region (2 kb upstream and 0.5 kb downstream of the TSS) (Fig. 7D). We next designed two pairs of chromatin immunoprecipitation (ChIP)–qPCR primers based on the sequences predicted with JASPAR and the open chromatin regions identified by ATAC-seq (Fig. 7E). ChIP assays with an anti-c-Myc antibody in KYSE150 and KYSE450 cells indicated that the VAMP3 promoter region was substantially immunoprecipitated (Supplementary Fig. S7A, B). Furthermore, c-Myc knockdown and overexpression positively regulated VAMP3 mRNA and protein levels in ESCC cells (Supplementary Fig. S7C-F).

**Fig. 7.**
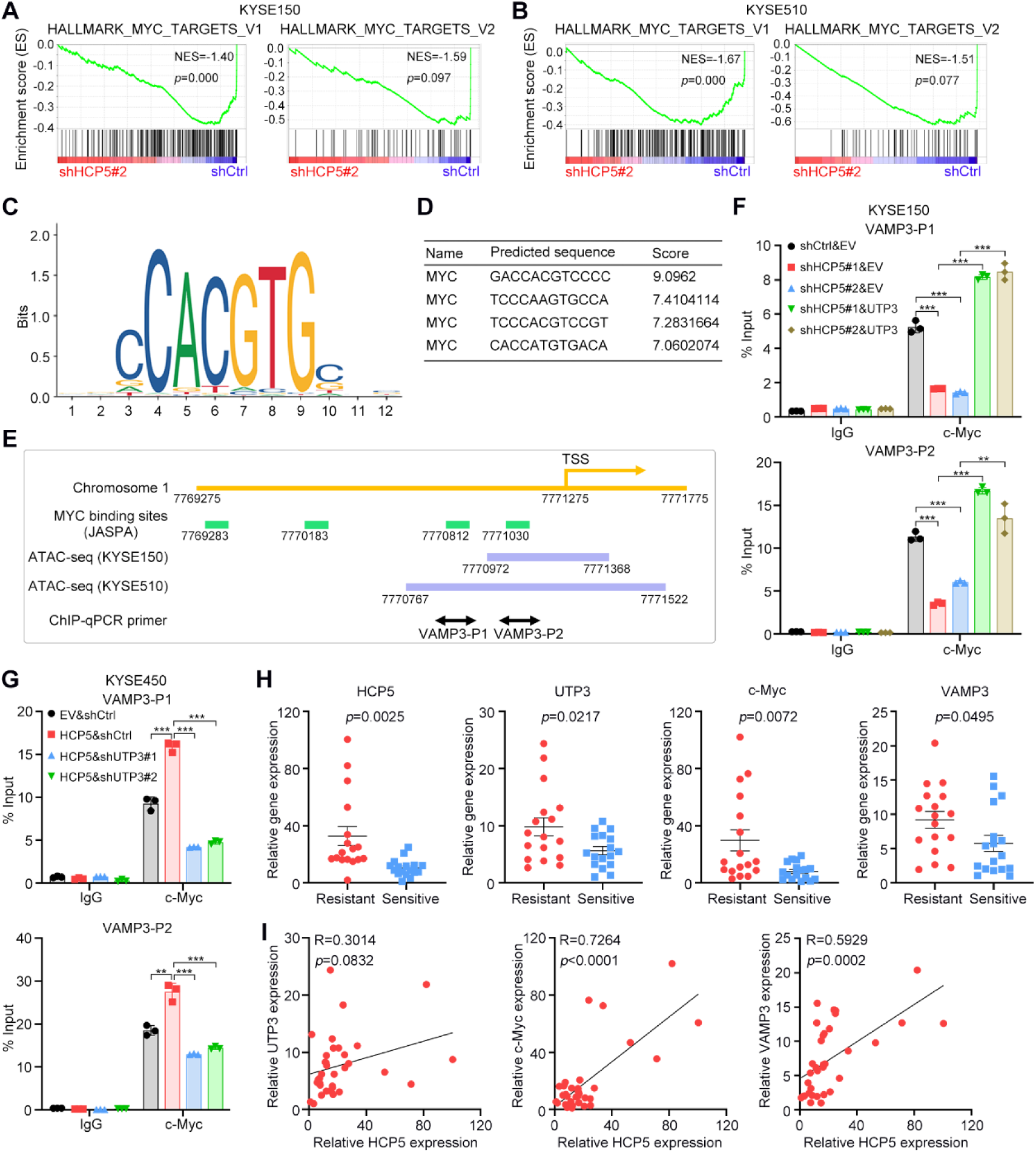
UTP3 recruits c-Myc to activate VAMP3 transcription. **A, B** GSEA results based on RNA-seq data comparing HCP5-silenced to control KYSE150 (A) and KYSE510 (B) cells. **C** Prediction of the sequence logo of the MYC-binding site based on the JASPAR database. **D** Predicted binding sequence of MYC in the VAMP3 promoter region, as determined from the JASPAR database. **E** Schematic showing MYC binding sites at the VAMP3 promoter region and the unique peaks identified by ATAC-seq. **F, G** ChIP analyses of the indicated cells using antibodies against c-Myc to identify the indicated gene promoter segments. The data are presented as the means ± s.d.; two-tailed *t* test, \*\*\**P* < 0.001, \*\**P* < 0.01; n = 3. **H** RT–qPCR detection of the expression of the indicated genes in samples from ESCC patients with chemoresistance or chemosensitivity. The data are presented as the means ± s.e.m.; two-tailed *t* test; n = 17 for chemoresistant patients and n = 17 for chemosensitive patients. **I** Correlation analyses between the indicated gene expressions in ESCC samples. Spearman correlation coefficients are shown.

To further investigate how HCP5/UTP3 affects the transcriptional regulation of VAMP3 by c-Myc, we first determined whether HCP5 affects c-Myc directly. No obvious changes in c-Myc expression levels were observed after HCP5 overexpression or knockdown (Supplementary Fig. S7G, H). There was also no direct interaction between HCP5 and c-Myc (Supplementary Fig. S7I, J). We then asked if HCP5 indirectly regulates c-Myc via UTP3. Co-IP assays confirmed the interaction between endogenous UTP3 and c-Myc (Supplementary Fig. S7K, L). Interestingly, ChIP–qPCR analyses indicated that UTP3 could also bind to the promoter region of VAMP3 (Supplementary Fig. S7 M, N). These results indicate that UTP3 may be a transcriptional coactivator that facilitates the recruitment of c-Myc to the VAMP3 promoter region. Supporting this speculation, a ChIP–qPCR assay showed decreased binding of c-Myc to the VAMP3 promoter region after HCP5 silencing, whereas the overexpression of UTP3 led to the complete rescue of c-Myc binding to the VAMP3 promoter (Fig. 7F). On the other hand, silencing UTP3 reversed the increased c-Myc binding to the VAMP3 promoter mediated by HCP5 overexpression (Fig. 7G).

Finally, we validated the clinical relevance of the HCP5/UTP3/c-Myc/VAMP3 axis in the baseline samples collected by pinch biopsy of 34 patients who received neoadjuvant chemotherapy (17 patients with chemosensitivity and 17 patients with chemoresistance). Notably, the expression levels of HCP5, UTP3, c-Myc and VAMP3 were significantly higher in the chemoresistant group than in the chemosensitive group (Fig. 7H). Further analysis showed that the expression of HCP5 was positively correlated with the expression of UTP3, c-Myc and VAMP3 in ESCC samples (Fig. 7I). These data demonstrated that the HCP5/UTP3/c-Myc/VAMP3 axis could serve as both predicative markers and therapeutic targets during chemotherapies of ESCC and potentially other cancer types.

## Discussion

Neoadjuvant chemotherapy is the first-line treatment strategy for most solid tumors, whereas drug resistance is a major cause of treatment failure and the primary contributor to the high mortality rate (35,36). Apoptosis resistance is a hallmark of cancer and is important for cancer progression and therapy resistance (37). The identification of novel anti- and proapoptotic molecules is essential for overcoming resistance to chemotherapy, radiotherapy or immunotherapy (38). To resolve drug resistance acquisition, we generated two DDP-resistant ESCC cell lines and performed whole-transcriptome sequencing in the resistant and parental cell lines (22). We identified HCP5 as the most significantly upregulated ncRNA in DDP-resistant cells and validated its function in preventing cancer cell apoptosis and contributing to chemoresistance. To date, HCP5 has been reported to be involved in the pathogenesis of multiple cancers, including driving fatty acid oxidation to promote chemoresistance (21) and facilitating malignant progression as a miRNA sponge (23–25). It has also been reported that HCP5 exerts a negative effect on cancer cell apoptosis (25,39); however, the potential mechanism for this remains unclear. We found that HCP5 inhibited caspase-dependent cancer cell apoptosis, and this effect was mediated by its interaction with UTP3, which prevents UTP3 from ubiquitination-mediated degradation.

UTP3 is a component of the SSU processome complex involved in ribosomal RNA (rRNA) processing and ribosome assembly (40) and mediates the derepression of silenced genes (26). These findings suggest that UTP3 may play an important role in gene transcriptional regulation. However, the mystery of UTP3 has not been unveiled, especially in the human context. Both the regulatory mechanism of UTP3 protein stability and its mediated downstream gene transcription remain unclear. Herein, we found that HCP5 interacted with and stabilized UTP3 by blocking the degradation of UTP3 induced by TRIM29-mediated ubiquitination. UTP3 then functions as a transcriptional coactivator to recruit c-Myc and subsequently activates VAMP3 transcription, which eventually leads to apoptosis resistance in cancer cells. Thus, our study filled the gap in knowledge of UTP3-mediated gene transcription and its stability regulation, raising the importance of further investigating the functions and mechanism of UTP3 during cancer development and progression.

c-Myc is a well-recognized oncoprotein that regulates cancer apoptosis and stemness (41,42). Although an increasing number of studies have revealed the regulatory network of c-Myc during cancer progression, the detailed mechanisms in specific contexts remain unclear. Our previous study revealed an essential role for E2F1/c-Myc signaling in the regulation of cancer stemness and chemoresistance of SCCs (43). Herein, we demonstrated HCP5/UTP3 as another upstream regulator of c-Myc-mediated cancer cell apoptosis resistance and chemoresistance. In addition, the results that VAMP3 functions as a critical downstream target of the HCP5/UTP3/c-Myc axis in the above phenotypes indicate the need to further explore the detailed mechanism underlying how VAMP3 regulates cancer cell apoptosis in future studies. Particularly, since VAMP3 is localized in recycling endosomes and regulates cytokine release (44), it would be potentially interesting to further investigate how the HCP5/UTP3/c-Myc/VAMP3 axis affects the tumor microenvironment that contributes to oncogenic roles.

## Materials and methods

### Cell lines and cell culture

The human ESCC cell lines were kindly provided by Dr. Yutaka Shimada (Kyoto University, Kyoto, Japan). The 293T cell line was purchased from the American Type Culture Collection (ATCC, Manassas, VA, USA). ESCC cells were maintained in RPMI 1640 plus 10% fetal bovine serum (FBS), and 293T cells were cultured in Dulbecco’s modified Eagle’s medium (DMEM) supplemented with 10% FBS. All cells were maintained in a humidified atmosphere with 5% CO_2_ at 37 °C. All cell lines were routinely authenticated using short tandem repeat (STR) DNA fingerprinting, tested for mycoplasma contamination with a MycoBlue Mycoplasma Detector (Vazyme Biotech, Nanjing, Jiangsu, China) and found to be negative for mycoplasma before use in any experiments.

### Antibodies and reagents

Antibodies against the following proteins were used for western blotting: SAS10 (1:5000, BETHYL Laboratories, Montgomery, TX, USA, A302-067A), cleaved Caspase-3 (1:1000, Cell Signaling Technology, MA, USA, #9664), cleaved Caspase-7 (1:1000, Cell Signaling Technology, #8438), cleaved PARP (1:1000, Cell Signaling Technology, #5625), MCL1 (1:1000, Cell Signaling Technology, #94296), c-Myc (1:1000, Cell Signaling Technology, #94296), TRIM29 (1:1000, Cell Signaling Technology, #5182), VAMP3 (1:2000, Abcam, Cambridge, UK, ab200657), ubiquitin (1:1000, Cell Signaling Technology, #3936), H3 (1:10000, Abcam, ab1791), GAPDH (1:5000, Sigma–Aldrich, St. Louis, MO, USA, G9545), and β-actin (1:5000, Sigma–Aldrich, A5316). Antibodies against the following proteins were used for IP: anti-SAS10 (1:100, BETHYL Laboratories, A302-067A), anti-TRIM29 (1:100, Cell Signaling Technology, #5182), anti-c-Myc (1:50, Cell Signaling Technology, #9402), and normal rabbit IgG (Cell Signaling Technology, #2729). SAS10 antibody (BETHYL Laboratories, A302-067A) was used for the RIP assay, and 5 μg was used in each IP reaction. The following antibodies were used for ChIP: anti-SAS10 (1:50, BETHYL laboratories, A302-067A) and anti-c-Myc (1:50, Cell Signaling Technology, #9402). The following antibodies were used for PLA and IF assays: anti-SAS10 (1:200, Abnova, Taipei, Taiwan, China, H00057050-B01P) and anti-TRIM29 (1:200, Affinity Biosciences, Cincinnati, OH, USA, DF8050). CHX (HY-12320) and Z-VAD-FMK (HY-16658B) were purchased from MedChemExpress (MCE, Lexington, MA, USA). MG132 (T2154) was purchased from TargetMol (Boston, MA, USA).

### Virus production and cell infection

Lentivirus was produced using 293T cells with the second-generation packaging system psPAX2 (#12260, Addgene, Watertown, MA, USA) and PMD2. G (#12259, Addgene) plasmids. Transfection was performed using Hieff Tranns Liposomal Transfection Reagent (40802, Yeasen, Shanghai, China) according to the manufacturer’s protocol. Cells were infected with lentivirus supplemented with 8 μg/ml polybrene (Sigma–Aldrich) twice within 48 h and then selected with 1 μg/ml puromycin (Sangon Biotech, Shanghai, China) or 200 μg/ml G418 (Sigma–Aldrich) for 10 days. The shRNA sequences can be found in Supplementary Table S1

### CCK-8 cell proliferation and cell viability assays

Cell viability was evaluated using a Cell Counting Kit-8 (CCK-8) kit (CK04, Dojindo Laboratories, Kumamoto, Japan) and was measured at an optical density (OD) of 450 nm with a Multiskan FC microplate photometer (Thermo Fisher Scientific, Waltham, MA, USA). For the cell proliferation assay, 2×10^3^ ESCC cells were seeded into 96-well microplates, and one plate was measured each day for 5 consecutive days. The relative cell proliferation rate was calculated by normalizing the absorbance read at 450 nm on days 2-5 to the absorbance measured on the first day. For the cell viability assay, 1×10^4^ ESCC cells were seeded into 96-well microplates, and drugs were added after cell adhesion. Cell viability was detected 24 h later according to the instructions of the CCK-8 kit manufacturer. Cell viability was calculated by normalizing the absorbance read at 450 nm for the experimental groups to that of the negative control group.

### Flow cytometry

Flow cytometry was performed using an apoptosis detection kit (AD10, Dojindo Laboratories) following the manufacturer’s instructions as previously described (45).

### Caspase 3/7 activity assay

Caspase 3/7 activity was measured using a luminescence Caspase Glo 3/7 assay kit (G8090; Promega, Madison, WI, USA) according to the manufacturer’s instructions. ESCC cells were pretreated with DDP at different concentrations for 24 h before measurement.

### Real-time quantitative PCR (RT–qPCR)

Total RNA extraction, mRNA reverse transcription and PCR analysis were performed as described previously (46). The lncRNA was reverse transcribed into cDNA using a lnRcute lncRNA First-Strand cDNA kit (4992908; Tiangen Biotech, Beijing, China) and quantified using a lnRcute lncRNA qPCR kit (4992886; Tiangen Biotech) according to the manufacturer’s instructions. The RT–qPCR primers are shown in Supplementary Table S2.

### Immunoprecipitation (IP) and western blotting

IP and western blotting were performed as previously described (46). Proteins for co-IP and ubiquitination assays were extracted using Cell Lysis Buffer for Western and IP (P0013; Beyotime Biotechnology, Shanghai, China). Nuclear proteins were extracted from cells using the ExKine Nuclear and Cytoplasmic Protein Extraction Kit (KTP3001; Abbkine, Wuhan, Hubei, China) according to the manufacturer’s protocol. In other cases, proteins were extracted using RIPA Lysis Buffer (CW2333; CWBIO, Taizhou, Jiangsu, China). The samples were separated on 10%–15% gels depending on the molecular weights of the proteins, and then the proteins were transferred onto polyvinylidene difluoride membranes (Merck Millipore, Billerica, MA, USA).

### RNA fluorescence in situ hybridization (FISH) and immunofluorescence (IF) staining

RNA FISH assays were performed using a Ribo fluorescent in situ hybridization kit (C10910, RiboBio, Guangzhou, China) according to the manufacturer’s instructions. A mix of probes targeting HCP5 was synthesized and labeled with Cy3 fluorescent dye. The nuclei were stained with Hoechst 33342 (Thermo Fisher Scientific). IF staining was performed as previously described (47). PLA was performed using a Duolink In Situ Red Starter Kit (DUO92101, Sigma–Aldrich) following the manufacturer’s protocol.

### Nuclear and cytoplasmic RNA fractionation analysis

Nuclear and cytosolic fractions were separated using a PARIS kit (AM1921, Thermo Fisher Scientific) according to the manufacturer’s instructions. The expression levels of HCP5 in the nucleus and cytoplasm of the ESCC cell lines were detected by RT–qPCR as described above. The expression level of HCP5 in the nucleus was normalized to U1 small nuclear RNA (snRNA), and in the cytoplasm, it was normalized to GAPDH.

### RNA pull-down assay and mass spectrometry (MS) analyses

HCP5 sense and antisense strands were synthesized by in vitro transcription with a TranscriptAid T7 high yield transcription kit (K0441, Thermo Fisher Scientific) according to the manufacturer’s instructions. Then, RNAs were purified using a GeneJET RNA purification kit (K0731, Thermo Fisher Scientific). Next, the RNAs were labeled with desthiobiotinylate using a Pierce RNA 3’ end desthiobiotinylation kit (20163, Thermo Scientific). An RNA–protein pull-down assay was performed using a Pierce Magnetic RNA–protein pull-down kit (20164, Thermo Fisher Scientific). In brief, 50 pmol labeled HCP5 and anti-HCP5 RNA were bound to streptavidin magnetic beads, and 200 µg of protein was used to bind the RNA. The RNA–protein bead mixture was incubated for 1 h at 4°C with rotation. Finally, the RNA-binding proteins were eluted, and a follow-up analysis was performed. Gel-based liquid chromatography-tandem MS experiments were performed by Beijing Qinglian Biotech Co., Ltd. The MS data were analyzed using MaxQuant software (version 1.5.3.30) and the UniProtKB/Swiss-Prot human database.

### RNA immunoprecipitation (RIP) assay

An RIP assay was performed using a Magna RIP RNA binding protein immunoprecipitation kit (17-700; Merck Millipore) according to the manufacturer’s instructions. Purified RNA was subjected to relative quantitation using RT–qPCR. The IP efficiency was calculated using the percent input method.

### CHX chase assay

The half-life of UTP3 was measured using a CHX chase assay. Cells were cultured for 24 h and treated with CHX (50 mg/mL) for 0, 12, 18 and 24 h. The cells were collected at the indicated time points, and total protein lysates were prepared for western blotting to detect UTP3. ImageJ software was used for the semiquantitative analysis of the UTP3 expression levels.

### In vivo ubiquitination assay

The in vivo ubiquitination assay was performed in ESCC cells. The cells were treated for 6 h with 20 mM MG132 before they were harvested. Protein in the cell lysate was immunoprecipitated to isolate ubiquitinated UTP3 with an anti-UTP3 antibody, then endogenous ubiquitin chains on UTP3 were detected.

### RNA-seq

The RNA-seq experiment and data analysis were performed by Seqhealth Technology Co., Ltd. (Wuhan, Hubei, China). Three replicates were prepared for each group using independent cell cultures. The paired-end libraries were prepared using a NEBNext Ultra RNA Library Prep Kit for Illumina (New England Biolabs, Ipswich, MA) following the manufacturer’s instructions, and sequencing was performed using the Illumina NovaSeq 6000 platform. A fold change of >2 and P value of <0.05 were set as the criteria for identifying significantly differentially expressed genes. The RNA-seq data can be accessed in the GEO database under accession number GSE200270.

### ATAC-seq

The ATAC-Seq experiment and data analysis were performed by Seqhealth Technology Co., Ltd.. In brief, 10 000–50 000 cells were treated with cell lysis buffer, and nuclei were collected after centrifugation for 5 min at 500 × g. Tagmentation was performed, and a high-throughput DNA sequencing library was prepared using TruePrep DNA Library Prep Kit V2 for Illumina (TD501; Vazyme Biotech). The library products were then enriched, quantified, and sequenced on a NovaSeq 6000 sequencer (Illumina, San Diego, CA) using the PE150 strategy. Raw data were filtered using Trimmomatic (version 0.36) and deduplicated with FastUniq (version 1.1). Clean reads were mapped to the reference genome with Bowtie2 software (version 2.2.6) and default parameters. RSeQC software (version 2.6) was used for read distribution analysis, and the insert length was calculated using Collect Insert Size Metrics tools in Picard software (version 2.8.2). Peak calling was performed with MACS2 software (version 2.1.1), and peak annotation and distribution analysis were performed using BEDtools (version 2.25.0). Differential peaks were identified using a Python script and Fisher’s exact test. The ATAC-seq data can be accessed in the Gene Expression Omnibus database under accession number GSE200373.

### ChIP Assay

ChIP assays were performed using a ChIP kit (#9003, Cell Signaling Technology) according to the manufacturer’s protocol. The ChIP qPCR primers targeting the VAMP3 promoter region were (5’ to 3’) VAMP3-P1-F, AGCAGATTATAATATGATC; VAMP3-P1-R, AGGGCACCATGTTTCTCTA; VAMP3-P2-F, GAGCCCAGAATAA-CATGAA; and VAMP3-P2-R, GGCGGTCCGGGCTCTGAAA.

### Xenograft assays

All animal experiments were approved by the Animal Care and Use Committee of the Chinese Academy of Medical Sciences Cancer Hospital. For subcutaneous xenografting, 1 × 10^6^ KYSE150 cells or 3 ×10^6^ KYSE450 cells were subcutaneously implanted into 6-week-old male BALB/c nude mice (HFK Bioscience, Beijing, China). One week after xenografting, mice were administered saline or DDP (6 mg/kg) every 3 days before being sacrificed. The PDX models were established with NOD-Prkdc^scid^-Il2rg^em1IDMO^ (NPI) mice (IDMO, Beijing, China) as previously described (47). AAV (AAV-shCtrl or AAV-shHCP5) produced by ViGene Biosciences (Jinan, Shandong, China) was intratumorally injected twice in three-day intervals with 2.5 × 10^11^ vg/mouse produced by. Tumor volumes were calculated using the following formula: 0.52×length×width^2^. CIs were calculated using the following formula: (value of DDP treated knockdown or overexpression group/value of DDP treated control group)/(value of saline treated knockdown or overexpression group/value of saline treated control group).

### Patient samples and RNA in situ hybridization (ISH)

For ISH, human ESCC tissue microarray slides were purchased from Servicebio (#G6040, Wuhan, Hubei, China). For RT–qPCR, 34 baseline samples were collected from patients receiving neoadjuvant chemotherapy followed by surgery at the Cancer Hospital, Chinese Academy of Medical Sciences. Written informed consent was obtained from all patients. RNA ISH was performed with an Enhanced Sensitive ISH Detection Kit I (POD) (MK1030; Boster, Wuhan, Hubei, China) according to the manufacturer’s instructions. Slides were scanned using an Aperio scanning system (Aperio, San Diego, CA, USA). H-scores were calculated as described previously (46). The sequences of the probes used for ISH were (5’ to 3’): Probe #1, GCAATGTAGTC-AACTCCTCACTTAGGGTCTGGTTGGTCAC; Probe #2, TAGGTAGCCTCATCC-CTGCCACTGTGTTTGCTCCATTTAT; and Probe #3, GCCCTCCACTGTGACTCT-CCTACTGGTGCTTGGTTCAGCT.

### Statistical analysis

Two-tailed Student’s t test, two-way ANOVA, or the Kaplan–Meier test was used for statistical analysis. For all statistical analyses, differences for which p ≤ 0.05 were considered statistically significant and results from at least three biologically independent experiments with similar results are reported. The data are presented as the means ± s.e.m. or means ± s.d. as indicated in the figure legends. The sample size (n) for each statistical analysis was determined on the basis of pretests and previous similar experiments and is indicated in the figure legends. Statistical analysis was performed with GraphPad Prism software (version 8.3.0, CA, USA).

## Acknowledgments

We are grateful to Dr. Yutaka Shimada (Kyoto University, Kyoto, Japan) for the ESCC cell lines. We thank Dr. Yin Li (Cancer Hospital, Chinese Academy of Medical Sciences, Beijing, China) for the ESCC samples. We appreciate Beijing Qinglian Biotech Co., Ltd., (Beijing, China) for performing the MS analysis. We thank Seqhealth Technology Co., Ltd. (Wuhan, Hubei, China) for performing the RNA-seq and ATAC-seq data analyses. Q.L. is supported by the Leukemia & Lymphoma Society.

## Author Contributions

Y.N. and Q.L. conceived this study. Y.N. performed most of the experiments. Y.N., Q.L. and X.W. analyzed the data. S.L., P.Z., and W.C. assisted with animal studies. Y.N. drafted the manuscript with assistance from Q.L. and Z.L. All authors reviewed the manuscript. Z.L. organized and supervised this study.

## Ethics approval and consent to participate

All animal protocols were approved by the Animal Care and Use Committee of the Chinese Academy of Medical Sciences Cancer Hospital.

## Data Availability Statement

The authors declare that the data supporting the findings of this study are available within the main text and supplementary materials. High-throughput sequencing data that support the findings of this study have been deposited in GEO database with the primary accession codes GSE200270 (RNA-seq) and GSE200373 (ATAC-seq).

