## Supplemental figures and tables for "HCP5 prevents ubiquitination-mediated UTP3 degradation to inhibit apoptosis by activating c-Myc transcriptional activity"

### Supplementary figures

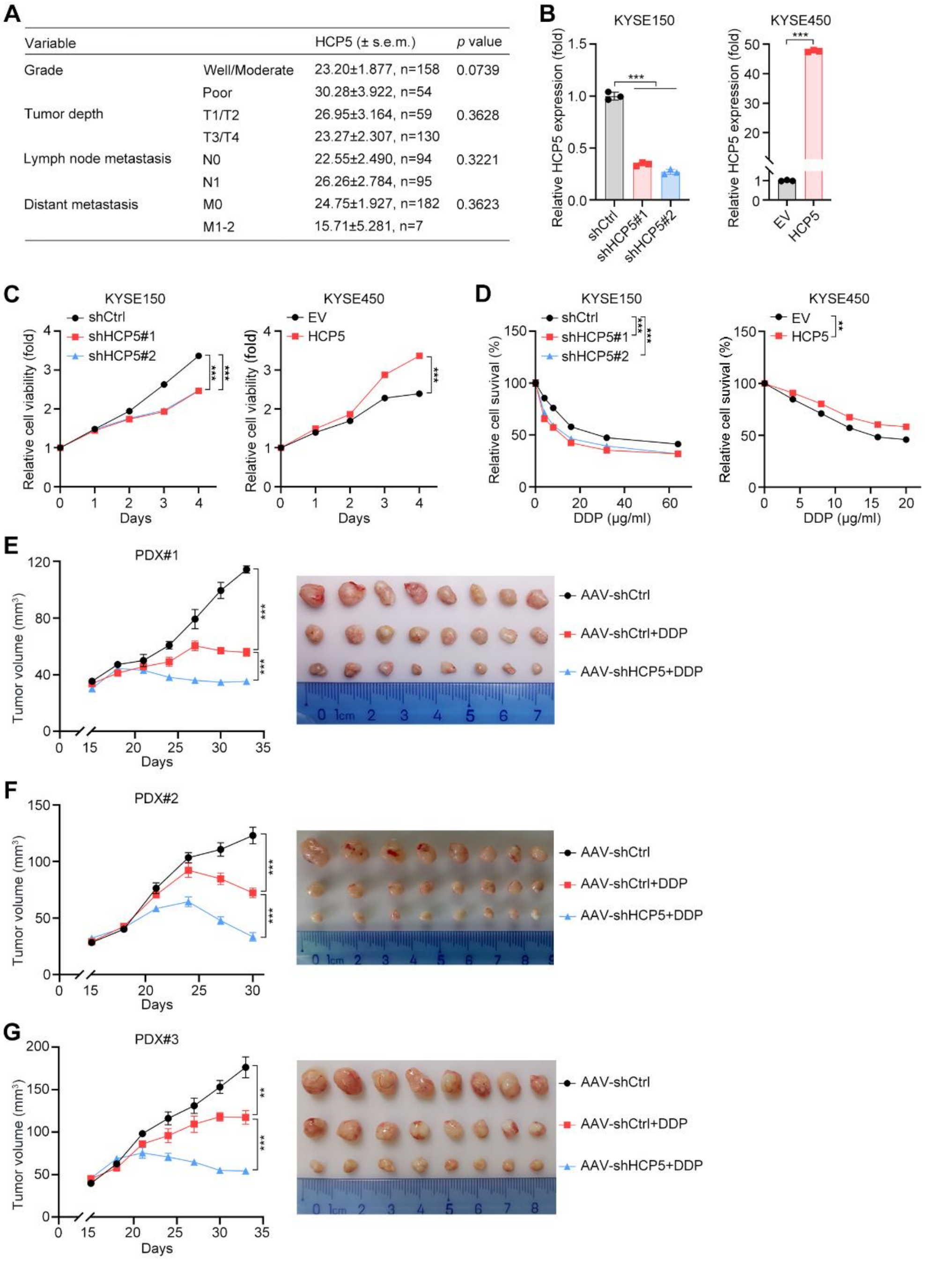


**Fig. S1 HCP5 promotes proliferation and chemoresistance in ESCC.**

**A** Correlations between HCP5 expression and clinical variables in patients with ESCC. Two-tailed *t* tests. **B** RT–qPCR detection of HCP5 expression in the indicated cells. The data are presented as the means ± s.d.; two-tailed *t* test, ***$P < 0.001$; n = 3. **C** In vitro growth curves of the indicated cells. The data are presented as the means ± s.d.; two-tailed *t* test, ***$P < 0.001$; n = 3. **D** Relative viability of the indicated cells after DDP treatment for 24 h. The data are presented as the means ± s.d.; two-way ANOVA, ***$P < 0.001$, **$P < 0.01$; n = 3. **E-G** In vivo growth curves and representative images of three PDX models treated with AAV-shCtrl or AAV-shHCP5. The data are presented as the means ± s.e.m.; two-tailed *t* test, ***$P < 0.001$, **$P < 0.01$; n = 8.

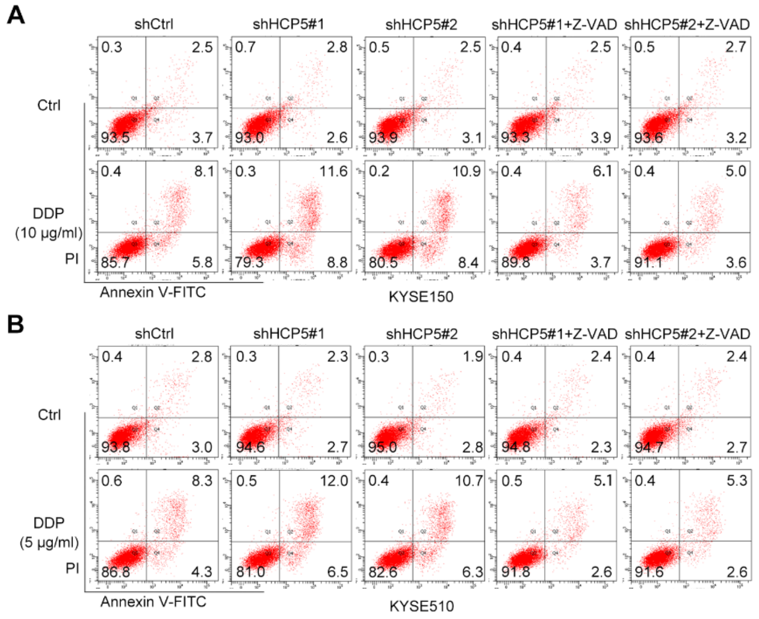


**Fig. S2 HCP5 impedes caspase-dependent apoptotic signaling.**

**A, B** Representative images of the flow cytometry detection of the apoptosis rate in the indicated cells treated with DDP with or without Z-VAD (50 μM).

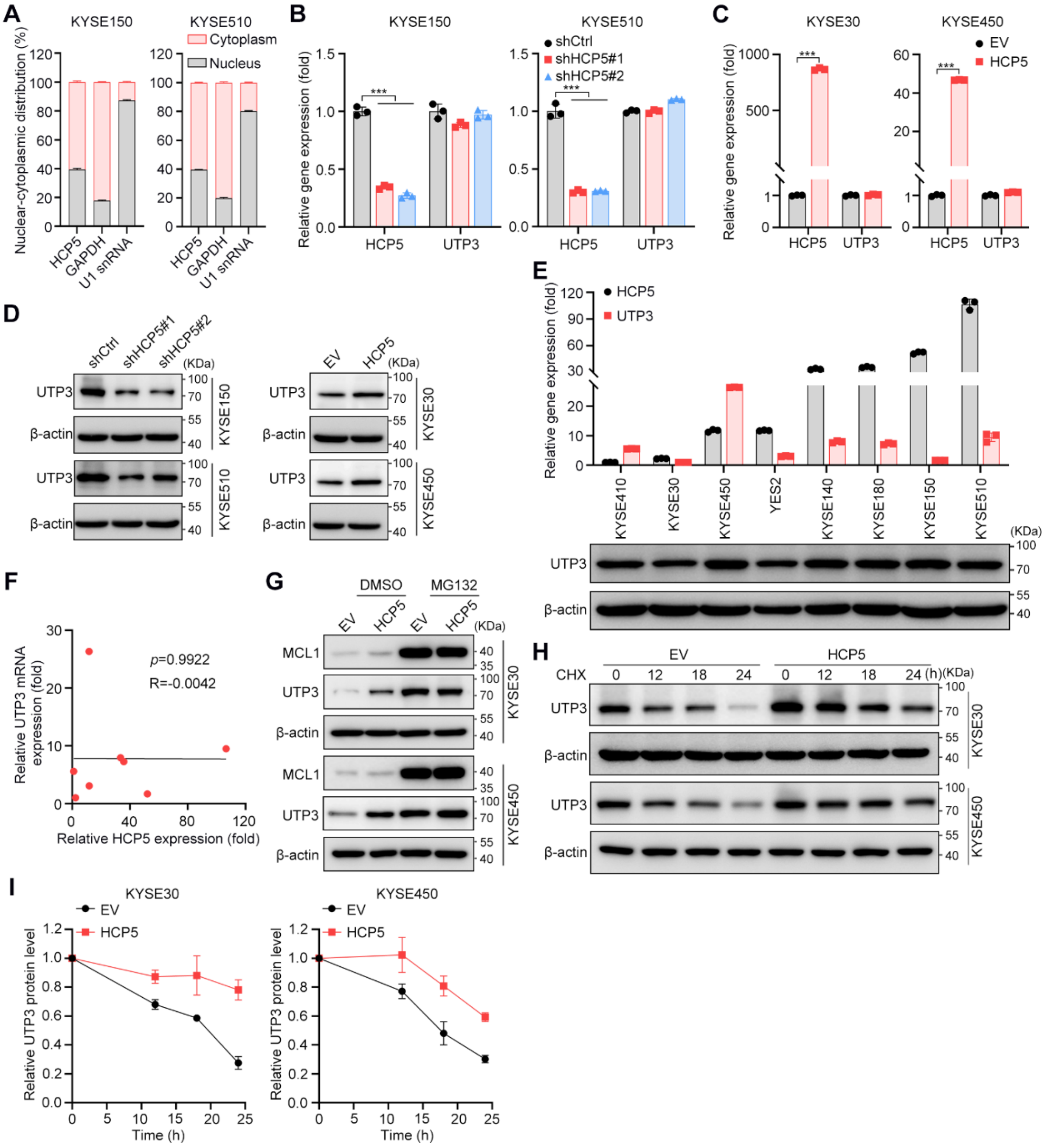


**Fig. S3 HCP5 stabilizes UTP3 by preventing its ubiquitination.**

**A** RT–qPCR detection of HCP5 expression in the cytoplasmic and nuclear fractions of KYSE150 and KYSE510 cells. **B, C** RT–qPCR detection of HCP5 and UTP3 expression in the indicated cells. The data are presented as the means ± s.d.; two-tailed *t* test, ****P* < 0.001; n = 3. **D** Western blot analysis of UTP3 expression in the indicated cells. **E** Western blot analysis of UTP3 protein expression and RT–qPCR detection of HCP5 and UTP3 mRNA expressions in ESCC cell lines. **F** Correlation analysis of

HCP5 and UTP3 mRNA expressions in ESCC cell lines. Spearman correlation coefficients are shown. **G** Western blot analyses of UTP3 and MCL1 expressions in KYSE30 and KYSE450 cells expressing EV or HCP5 treated with DMSO or MG132 for 8 h. **H** Western blotting analyses of UTP3 expression in KYSE30 and KYSE450 cells expressing EV or HCP5 subjected to a CHX pulse-chase assay. **I** Quantitative analyses of UTP3 expression in KYSE30 and KYSE450 cells expressing EV or HCP5 subjected to a CHX pulse-chase assay. The quantification of UTP3 expression relative to $\beta$-actin expression is shown. The data are reported as the means ± s.e.m.; n = 3.

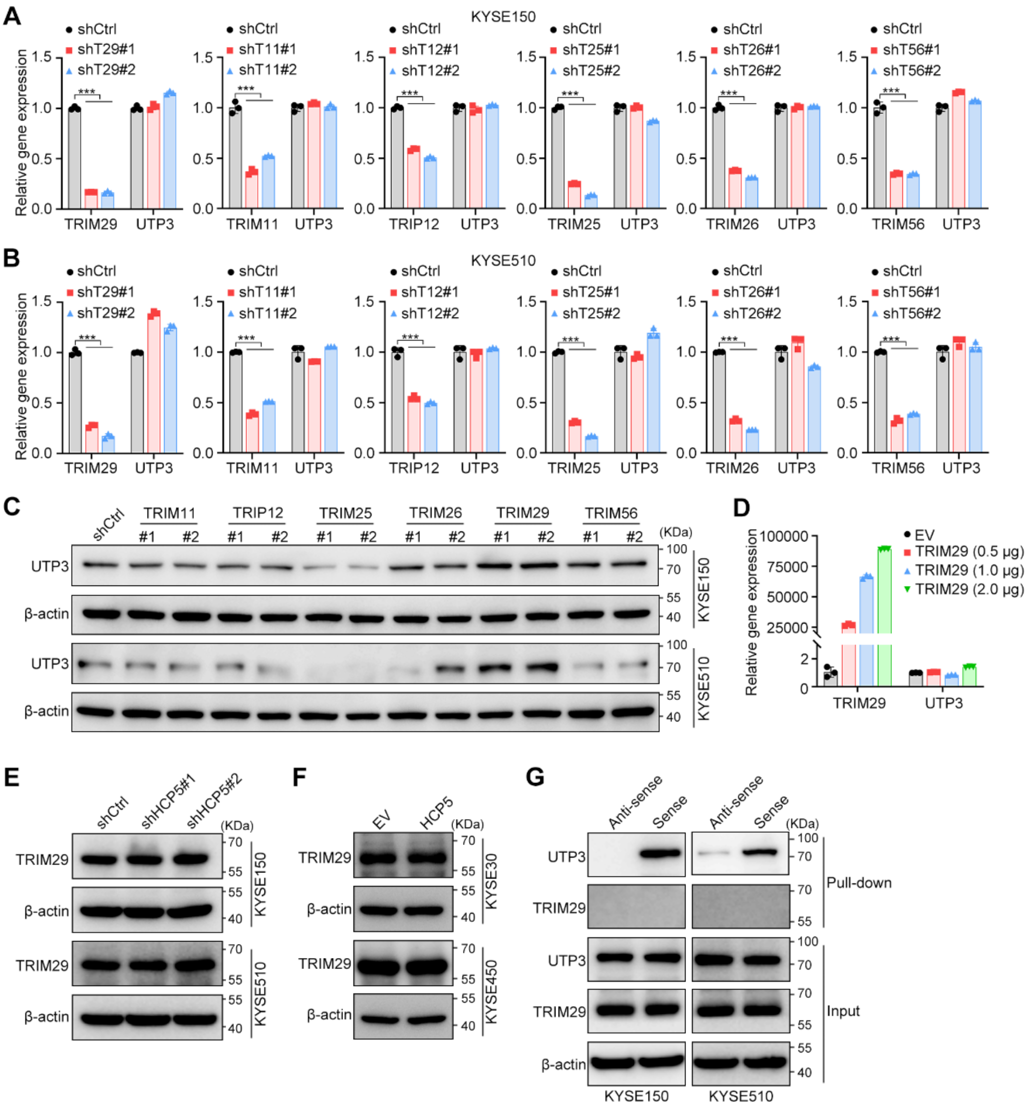


**Fig. S4 TRIM29 interacts with and ubiquitinates UTP3.**

**A, B** RT–qPCR measurement of the UTP3 and E3 ligases expression levels in the indicated cells. The data are presented as the means ± s.d.; ***$P$ < 0.001; n = 3. **C** Western blot analysis of UTP3 expression in the indicated cells. **D** RT–qPCR measurement of TRIM29 and UTP3 expressions in 293T cells transfected with increasing amounts of TRIM29. **E, F** Western blot analysis of TRIM29 expression in the indicated cells. **G** Western blot analyses of TRIM29 and UTP3 in precipitates from RNA pull-down assays performed using antisense or sense sequences of HCP5.

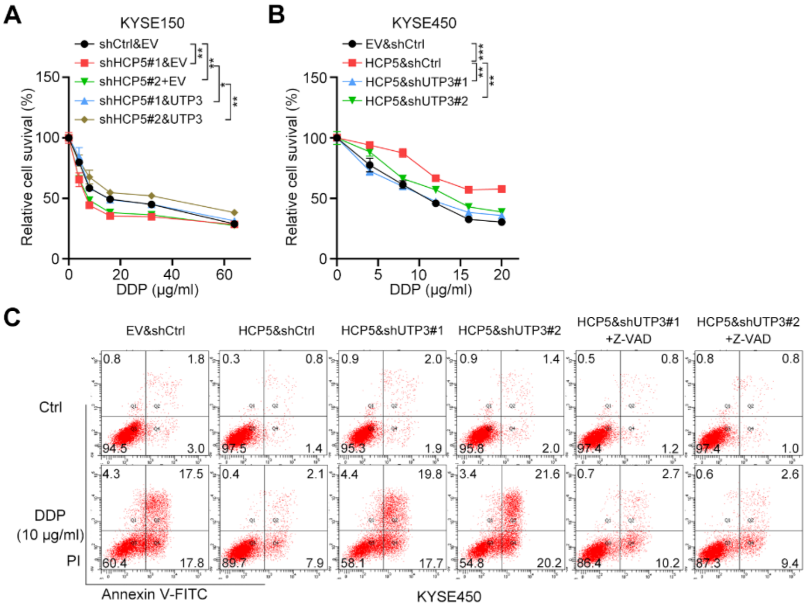


**Fig. S5 HCP5 inhibits caspase-dependent cell apoptosis via UTP3.**

**A, B** Relative viability of the indicated cells after DDP treatment for 24 h. The data are presented as the means ± s.d.; two-way ANOVA, ***$P < 0.001$, **$P < 0.01$, *$P < 0.05$; n = 3. **C** Representative images of the flow cytometry detection of the apoptosis rate in the indicated cells treated with DDP with or without Z-VAD (50 μM).

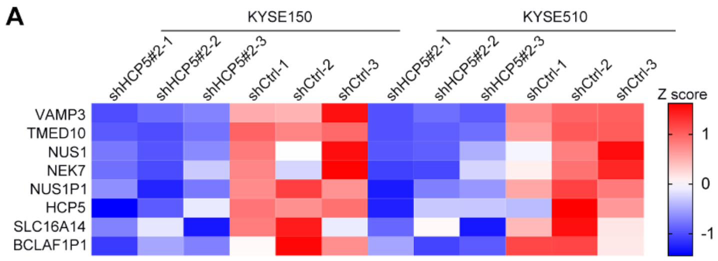


**Fig. S6 HCP5 activates VAMP3.**

**A** Heatmap showing the expression profile of the 8 consistently downregulated genes based on RNA-seq data obtained from HCP5-silenced KYSE150 and KYSE510 cells.

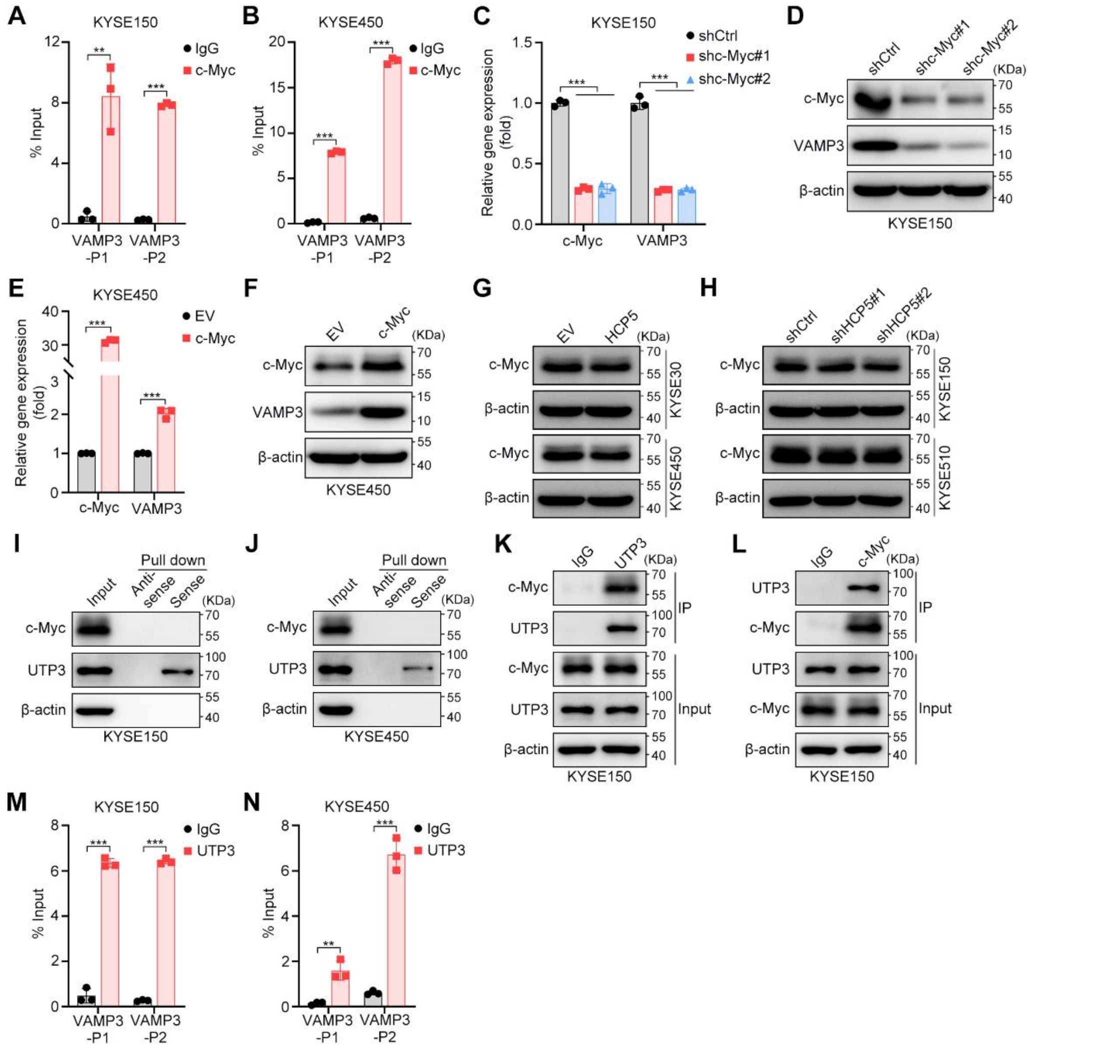


**Fig. S7 UTP3 recruits c-Myc to activate VAMP3.**

**A, B** ChIP assay of the VAMP3 promoter region in KYSE150 (A) and KYSE450 (B) cells with antibodies against c-Myc. The data are presented as the means ± s.d.; two-tailed *t* test, ****P* < 0.001, ***P* < 0.01; n = 3. **C-F** RT–qPCR and Western blot analyses of the expression of c-Myc and VAMP3 in the indicated cells. The data are presented as the means ± s.d.; two-tailed *t* test, ****P* < 0.001; n = 3. **G, H** Western blot analyses of c-Myc expression in the indicated cells. **I, J** Western blot analyses of c-Myc and UTP3 in the precipitates from RNA pull-down assays performed using HCP5 antisense or sense sequences. **K, L** Western blot analyses of c-Myc and UTP3 in precipitates from co-IP assays with KYSE150 cells. **M, N** ChIP assay of the VAMP3 promoter region in KYSE150 and KYSE450 cells with antibodies against UTP3. The data are presented as the means ± s.d.; two-tailed *t* test, ***$P$ < 0.001, **$P$ < 0.01; n = 3.

**Supplementary tables**

**Table S1. shRNA sequences**

| Sequence (5’ to 3’) | Name |
|---|---|
| GCGTTGAGCTGTGCCTATAGA | shHCP5#1 |
| GGTCTGGTTGGTCACCTAAAG | shHCP5#2 |
| GCTGTTACAGATCTTTCTGAT | shUTP3#1 |
| CCTAGGAGAAAGAAGATTGAT | shUTP3#2 |
| CGACAAGAACTCCAACTACTT | shTRIM29#1 |
| ACCTGAGCCGTAACTTCATTG | shTRIM29#2 |
| GGAAACGACGAGAACAGTTGA | shc-Myc#1 |
| CCTGAGACAGATCAGCAACAA | shc-Myc#2 |
| GCAGCCAAGTTGAAGAGGAAA | shVAMP3#1 |
| TGGTGGACATAATGCGAGTTA | shVAMP3#2 |
| GCACGGCTCTATCTCATCAAC | shTRIM56#1 |
| GCAGCAGAATAGTGTGGTAAT | shTRIM56#2 |
| GACACCGAGAGAAGCTGCACTACTA | shTRIM26#1 |
| CGAGAGAAGCTGCACTACTACTGTG | shTRIM26#2 |
| GCATCTGCTACGGAAGCATGA | shTRIM25#1 |
| GGTGGAGCAGCTACAACAAGA | shTRIM25#2 |
| CAAAGTGCAGTCTAATACTACTTCT | shTRIP12#1 |
| AGTGCAGTCTAATACTACTTCTGAA | shTRIP12#2 |
| GGTCACTGCTATTCATCTTTC | shTRIM11#1 |
| GGAGAACGTGAACAGGAAGGA | shTRIM11#2 |

**Table S2. RT–qPCR primers**

| Sequence (5' to 3') | Name |
|---|---|
| CCGGGAAACTGTGGCGTGATGG | GAPDH-F |
| AGGTGGAGGAGTGGGTGTCGCTGTT | GAPDH-R |
| AGTTGATCAAGGCCGTGGAA | HCP5-F |
| TTCCTGGGACACAGAGGACT | HCP5-R |
| GCCCTAGATATGGACGATGAGG | UTP3-F |
| CTCTGACCCCACGACAAACTG | UTP3-R |
| GTGGTGGACATAATGCGAGTT | VAMP3-F |
| CGTCTGCACGGTCGTCTAAC | VAMP3-R |
| AGGGAGATCCGGAGCGAATA | c-MYC-F |
| GTCCTTGCTCGGGTGTTGTA | c-MYC-R |
| AGGATGCCAAGACCACCAAC | TRIM29-F |
| TTGCAATGACAGCTCCGTCT | TRIM29-R |
| GAGAACGTGAACAGGAAGGAG | TRIM11-F |
| CCATCGGTGGCACTGTAGAA | TRIM11-R |
| ATGTCCAACCGGCCTAATAACA | TRIP12-F |
| TCCTATTGAGTCGTCTTGTGGT | TRIP12-R |
| AATCGGCTGCGGGAATTTTTC | TRIM25-F |
| TCTCACATCATCCAGTGCTCT | TRIM25-R |
| TGCACTACTACTGTGAGGACG | TRIM26-F |
| TCCTTAGGGTACTCAGGTGGT | TRIM26-R |
| GCCTGCATACCTACTGCCAAG | TRIM56-F |
| GCAGCCCATTGACGAAGAAGT | TRIM56-R |
| CCACTGGGGTGGTAAAACTTG | NEK7-F |
| AAGGACTTTGTAATGCAGCCAT | NEK7-R |
| GACGGTCGTTCCTTGGAGAAG | NUS1-F |
| CCACGGCCATACACCACAC | NUS1-R |
| AGATTCACAAGGACCTGCTAGT | TMED10-F |
| TCATGTCTAGGATCACGAGTTGG | TMED10-R |
| GTGTCCTCAACGTGGAATGG | SLC16A14-F |
| ATGAACAAGCCGATGAAAGGG | SLC16A14-R |
